# Generating Artificial Sensations with Spinal Cord Stimulation in Primates and Rodents

**DOI:** 10.1101/2020.05.09.085647

**Authors:** Amol P. Yadav, Shuangyan Li, Max O. Krucoff, Mikhail A. Lebedev, Muhammad M. Abd-El-Barr, Miguel A.L. Nicolelis

## Abstract

For patients who have lost sensory function due to a neurological injury such as spinal cord injury (SCI), stroke, or amputation, spinal cord stimulation (SCS) may provide a mechanism for restoring somatic sensations via an intuitive, non-visual pathway. Inspired by this vision, here we trained rhesus monkeys and rats to detect and discriminate patterns of epidural SCS. Thereafter, we constructed psychometric curves describing the relationship between different SCS parameters and the animal’s ability to detect SCS and/or changes in its characteristics. We found that the stimulus detection threshold decreased with higher frequency, longer pulse-width, and increasing duration of SCS. Moreover, we found that monkeys were able to discriminate temporally- and spatially-varying patterns (i.e. variations in frequency and location) of SCS delivered through multiple electrodes. Additionally, sensory discrimination of SCS-induced sensations in rats obeyed Weber’s law of just noticeable differences. These findings suggest that by varying SCS intensity, temporal pattern, and location different sensory experiences can be evoked. As such, we posit that SCS can provide intuitive sensory feedback in neuroprosthetic devices.

## Main

Lack of sensory feedback from a brain-controlled actuator or prosthetic device is a major hindrance to successful integration of the neuroprosthesis in activities of daily life and rehabilitative protocols (1-3). The somatosensory cortex (S1) and thalamus have been proposed as potential targets for neurostimulation that could produce naturalistic somatosensory percepts (4-10). However, stimulating these brain areas requires surgical implantation of deep intracranial electrodes – a procedure associated with significant risks. While peripheral nerve stimulation provides a less invasive alternative, sensations evoked with this method are highly localized, and thus limited in their applicability as a general-purpose sensory input pathway to the brain (11-13). Previously, our group demonstrated that electrical stimulation of the dorsal surface of the spinal cord can be used to transmit sensory information to the brain or between multiple brains in rodents (14). Building on this previous work, here we explored whether rats and nonhuman primates can learn to detect and discriminate artificial sensations generated with dorsal thoracic epidural spinal cord stimulation (SCS). Understanding the psychophysical relationship between SCS parameters and the ability to detect sensations is critical for the development of novel neuroprosthetic devices that restore function to people with sensory disabilities. We examined how sensory discrimination changes when SCS parameters are varied in both rodent and primate models, and we asked whether animals can learn to discriminate sensations generated by SCS patterns that vary in frequency and spatial location. After training the animals to discriminate SCS patterns, we determined whether artificial sensations evoked by SCS of variable frequency follow Weber’s law of just noticeable differences (JND) - a critical property defining sensory discrimination.

## Results

We implanted three rhesus monkeys with percutaneous epidural SCS electrodes at the dorsal thoracic spinal level and trained them to perform a two-alternative forced choice task (2AFC) using a joystick-controlled cursor (Figure 1a, Supplementary Figures 1b, 1c, and 1d). In a typical experimental session, a monkey was seated in a chair in front of a monitor that displayed task-related cues. The animals moved a hand-held joystick to control a cursor on a screen (Figure 1b).

**Figure 1:**
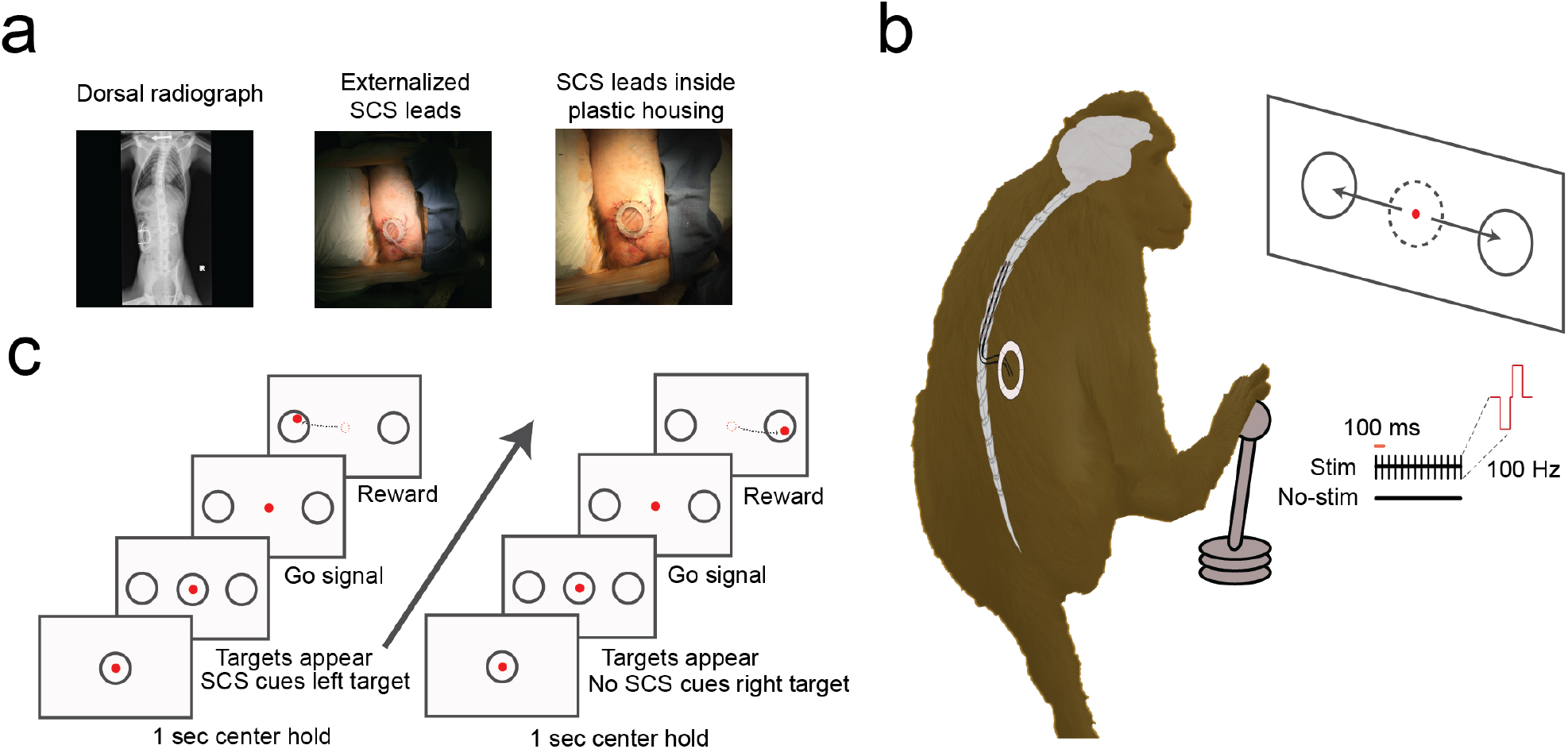
Surgery and experimental task setup. a) We implanted three non-human primates (rhesus monkeys) with SCS percutaneous leads over the T6-T10 dorsal epidural surface of the spinal cord. Leads were externalized from the lower back area and secured inside a custom plastic housing. Leads were manually accessed by the experimenter for daily training and connected to a custom pulse stimulator. b) Monkeys were seated in a primate chair in front of a computer monitor with access to a hand-controlled joystick. They participated in a two-alternative forced choice task (2AFC) by moving the joystick controlling a cursor on the screen in order to receive a juice reward. c) On each trial, monkeys had to hold the cursor inside the center circle for 1 sec. After that, targets appeared on the left and right side of the center. Monkeys were presented with ‘SCS-ON’ (biphasic, 100 Hz, 200 μs, 1 sec) cue or ‘SCS-OFF’ cue when the targets appeared. After a brief, variable hold period (100-1000 ms), the center circle disappeared which indicated them to move the joystick. Monkeys had to move the cursor inside the left target on ‘SCS-ON’ trials and inside the right target on ‘SCS-OFF’ trials. Correct response resulted in juice reward. In the SCS discrimination task, stimulation was delivered at 200 μs for 1 sec. Monkeys had to select left target for 100 Hz stimulus and right target for 200 Hz stimulus in the frequency discrimination task. For spatial discrimination task, monkeys had to choose left target when stimulation was delivered at electrode pair 1 and right target for stimulation at electrode pair 2.

A typical trial consisted of a brief (300 - 500 ms) center hold period after which two targets appeared. After a brief preparatory period (250 – 1000 ms) during which a trial cue was presented, the monkeys had to move the cursor into one of the targets to obtain a juice reward. Monkeys were initially trained to identify the correct target using a visual cue; however, during the experimental sessions, no visual cues were presented, and they selected a target by interpreting SCS cues alone. In the detection task, monkeys had to select the left target if SCS was delivered during the preparatory period and right target if no SCS was delivered (Figure 1c). In the discrimination task, monkeys had to select the left target for the 100 Hz stimuli and right target for the 200 Hz stimuli (frequency discrimination) and the left target for electrode pair 1 and right target for electrode pair 2 (spatial discrimination).

### Monkeys learned to detect SCS stimuli

Monkeys M, O, and K learned to detect SCS-induced sensory percepts evoked using percutaneous dorsal thoracic epidural electrodes (T7 for monkey M, T5-T6 for monkey O, T5-T6 for monkey K). Performance of all monkeys started below chance levels of 50% and reached above 90% after learning (Figure 2a). Monkey M started detection performance at 49% and reached a maximum of 93% in 16 days; monkey O started at 49% and reached a maximum of 97% in 10 days; and monkey K started at 41% and reached a maximum of 90% in 8 days.

**Figure 2:**
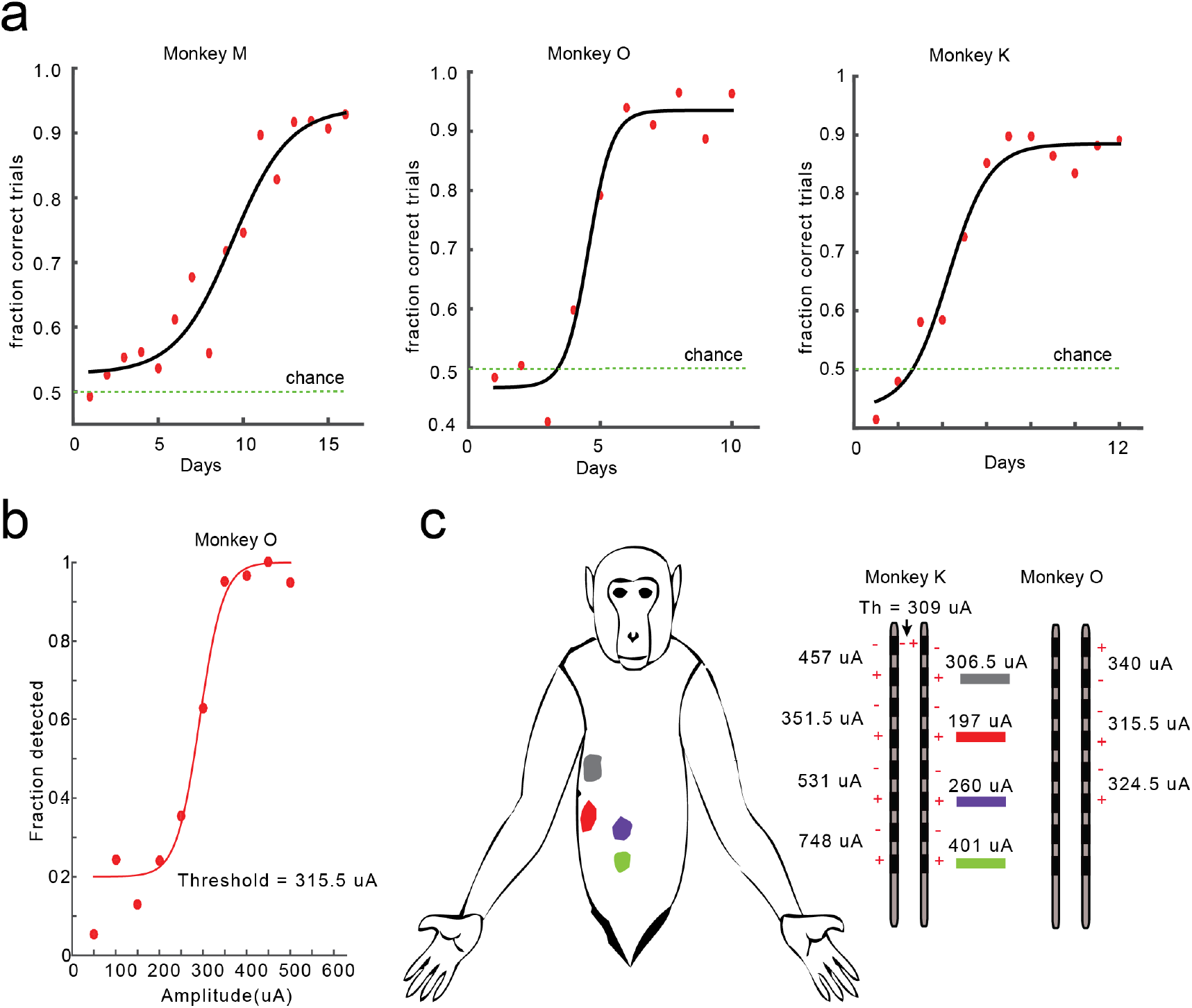
Monkeys learned to detect SCS stimuli. a) Learning curves (sigmoidal fits) for monkeys M, O, and K showing behavioral performance (fraction correct trials) as a function of training days. b) Psychometric function showing fraction trials detected as a function of stimulation amplitude in monkey O. Detection threshold is defined as amplitude at which monkeys achieved 75% performance on detection task. c) Mapping of bipolar electrode pairs on monkey K’s body where stimulation on right-side electrode at suprathreshold amplitude elicited minor muscle twitches or skin flutter (color coded by electrode pairs and corresponding sensory thresholds shown on right). Monkey body shape is adapted from (15).

### Electrode thresholds and electrode mapping

Once the monkeys learned to detect SCS sensations, we used psychometric analysis to determine the detection thresholds for different electrode combinations (Figure 2b). We observed that the detection thresholds varied from 315.6 μA to 340 μA for monkey O and 197 μA to 748 μA for monkey K for different cathode-anode electrode pairs (Figure 2c, right). Once electrode thresholds were determined, we mapped the bipolar electrode pairs to locations on the monkey’s body by stimulating at suprathreshold amplitudes and observing stimulation-induced minor muscle twitches or skin flutter. We observed that muscle twitches/skin flutter were elicited in the trunk and abdomen area only at suprathreshold values but not at sensory threshold values (Figure 2c). We also noted that in both monkeys K and O experimentally determined sensory thresholds were always lower than the observed motor thresholds for each cathode-anode electrode pair (Supplementary Figure 3b).

### Sensitivity to detection of sensory percepts in primates

Thereafter, we investigated the psychophysical relationship between stimulation parameters and detection of sensory percepts by varying stimulation amplitudes along with stimulation frequency, pulse-width, or duration of stimulation while keeping the other two parameters constant.

We varied amplitude from 50 μA to 800 μA for pulse-widths of 50 μs, 100 μs, 200 μs, and 400 μs for monkey K, and pulse-widths of 100 μs, 200 μs, and 400 μs for monkey O. Frequency and duration of stimulation were held constant (Figures 3a, 3e, and Supplementary Figure 2a). We observed that stimulus detection threshold significantly decreased with increasing stimulation pulse-width (p<0.05, repeated measures one-way ANOVA) for both animals (Figure 3h).

**Figure 3:**
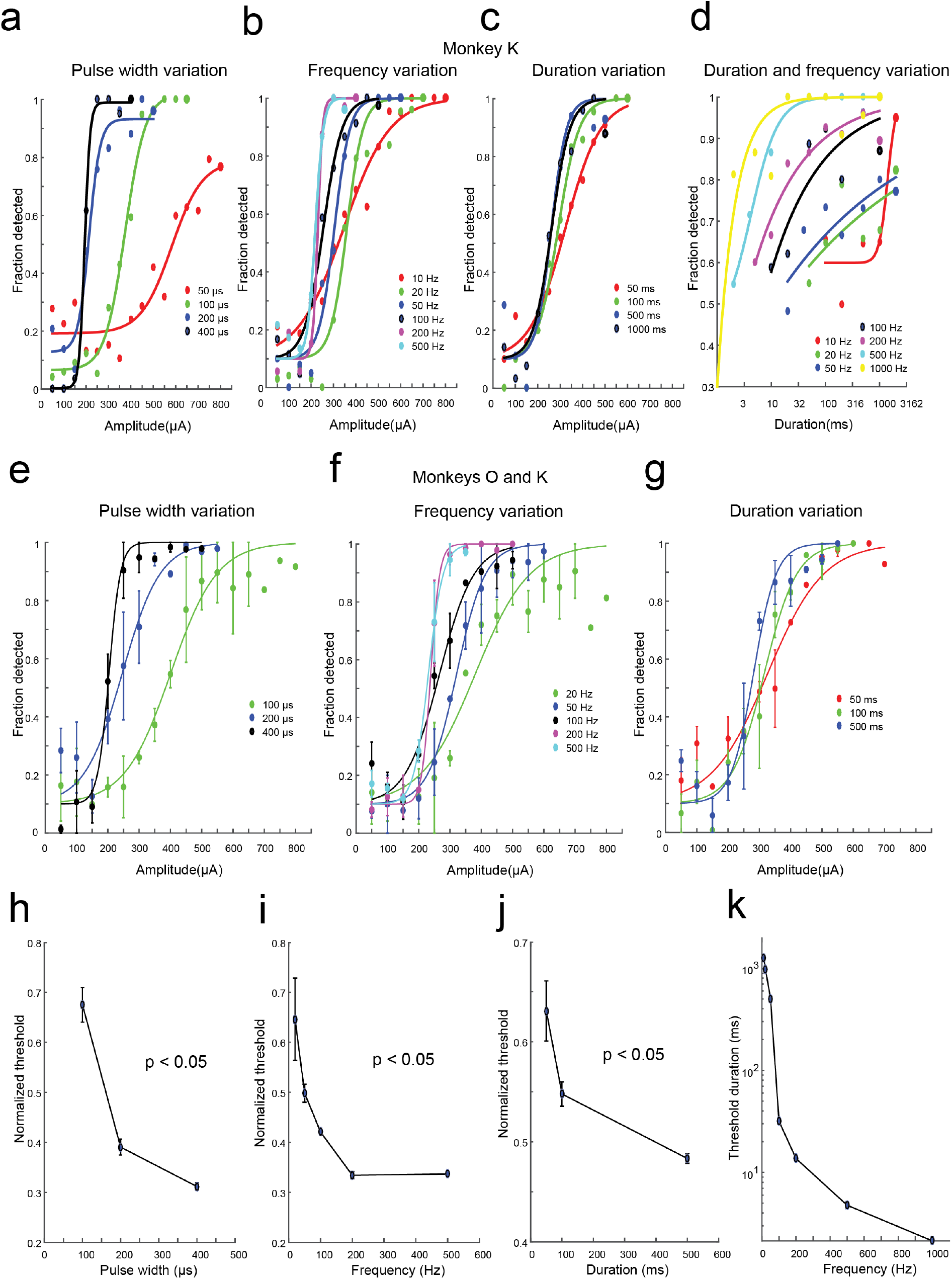
Psychophysical evaluation of SCS sensory detection with varying stimulation parameters in primates. Once monkeys learned to detect SCS stimuli at the standard parameters (frequency: 100 Hz, pulse width: 200 μs, duration: 1 sec), we allocated different blocks of sessions where (pulse width & amplitude; panels ‘a’ and ‘e’); (frequency & amplitude; panels ‘b’ and ‘f’); and (duration & amplitude; panels ‘c’ and ‘g’) were varied while keeping other parameters constant. In monkey K, in a separate block, frequency and duration was varied with other parameters constant (pulse width: 200 μsec and amplitude: 325 μA). Psychometric curves in a-g are sigmoidal fits. Panels a-d represent psychometric curves for monkey K, while panels e-g represent psychometric curves fitted to data averaged across monkeys K and O (circles and error bars are mean ± sem). Panels h, i, j, indicate normalized detection thresholds (normalized by maximum amplitude used in the experiment block of each individual monkey) averaged across both monkeys (mean ± sem). Detection threshold were calculated as 75% fraction detected at the stimulation parameters shown in panels a-c (monkey K) and Supplementary Figure 2 a-c (monkey O). P-values were calculated using repeated measures one-way ANOVA. Panel k shows threshold duration obtained as 75% detection from curves in panel d as a function of frequency.

We varied amplitude from 50 μA to 800 μA for frequencies of 10 Hz, 20 Hz, 50 Hz, 100 Hz, 200 Hz, and 500 Hz for monkey K, and frequencies of 20 Hz, 50 Hz, 100 Hz, 200 Hz, and 500 Hz for monkey O while keeping pulse-width and duration of stimulation constant (Figures 3b, 3f, and Supplementary Figure 2b). We observed that stimulation detection threshold significantly decreased with increasing stimulation frequency (p<0.05, repeated measures one-way ANOVA) for both animals (Figure 3i).

We varied amplitude from 50 μA to 600 μA for duration of 50 ms, 100 ms, 500 ms, and 1000 ms for monkey K, and amplitude between 50 μA to 700 μA for duration of 50 ms, 100 ms, and 500 ms for monkey O, while keeping pulse-width and frequency of stimulation constant (Figures 3c, 3g, and Supplementary Figure 2c). We observed that stimulation detection threshold significantly decreased with increasing stimulation duration (p<0.05, repeated measures one-way ANOVA) for both animals (Figure 3j).

In monkey K, we varied both frequency and duration of stimulation while keeping amplitude and pulse-width of stimulation constant. We observed that as the frequency of stimulation increased, the duration of stimulation to reach detection threshold decreased (Figures 3d and 3k). Monkey K was able to detect a sensory percept generated by merely two stimulation pulses delivered at 1000 Hz.

### Sensitivity to detection of sensory percepts in rats

We have previously shown that rats learn to detect sensations generated by epidural SCS delivered at the T2 spinal level (14). In order to study the psychophysical performance of rats pertaining to sensory detection, initially we trained rats to detect SCS stimuli using a 2-AFC task in a slightly modified behavioral chamber (Figure 4a and 4b). We trained five rats to detect SCS stimuli delivered at the T3 spinal level (frequency: 100 Hz, pulse-width: 200 μs, duration: 1 sec, biphasic pulses at 243.7±57.9 μA, Figure 4c).

**Figure 4:**
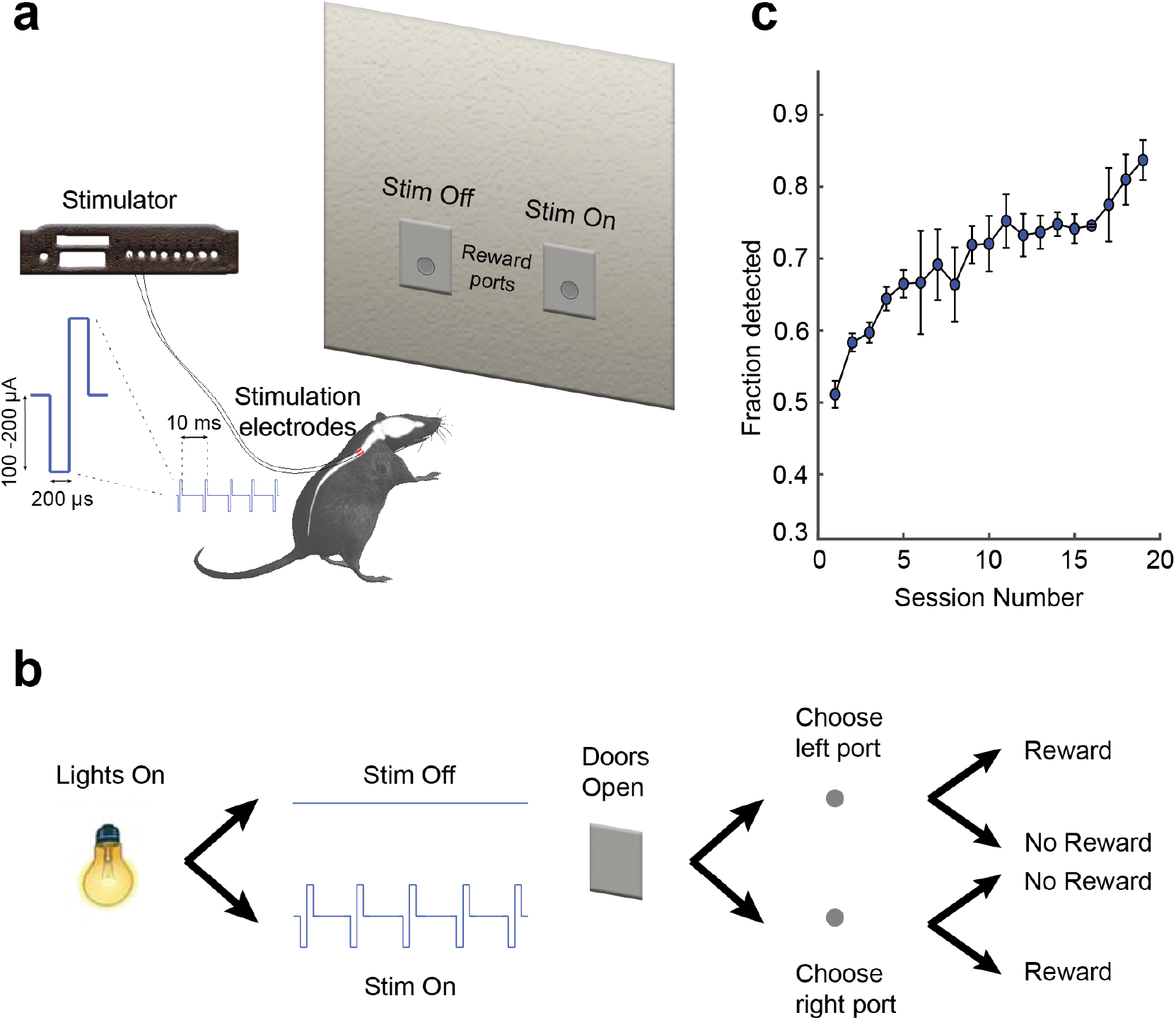
Behavioral task setup and rats learned to detect SCS stimuli. a) The experiment task consisted of a closed behavioral chamber with two reward ports on one side of the chamber. The ports were covered with vertical sliding doors. Five rats were implanted with bipolar stimulation electrodes on the dorsal surface of the spinal cord at the T3 spinal level. b) Task consisted of a house light turning ‘on’, followed by sensory cues for 1 second. During the cue period SCS was either delivered or not delivered. After the cue period both reward doors opened, and the rat had to make a nose poke response in either of the ports to receive water reward. If SCS was delivered rats had to poke inside the left port and if not delivered, then they had to poke inside the right reward port. Poking in the correct reward poke initiated a water reward, while incorrect pokes were not rewarded. c) Rats learned to detect SCS stimuli over a period of 15 – 25 days as indicated – learning curve showing task performance indicated by fraction trials detected as a function of training sessions. Circles and error bars indicate mean ± sem.

Once rats learned the basic detection task, we varied stimulation parameters such as: pulse width (50 – 400 μs); frequency (10 – 500 Hz); and duration (50 – 1000 ms) independently with stimulation amplitude (Figures 5a, 5d – pulse width; 5b, 5e – frequency; and 5c, 5f – duration). Similar to the results in monkeys, we observed that stimulation detection thresholds significantly decreased with increasing pulse-width (Figure 5g, p< 0.0001, repeated measures one-way ANOVA), frequency (Figure 5h, p< 0.0001, repeated measures one-way ANOVA) and duration (Figure 5i, p<0.05, repeated measures one-way ANOVA) for all rats.

**Figure 5:**
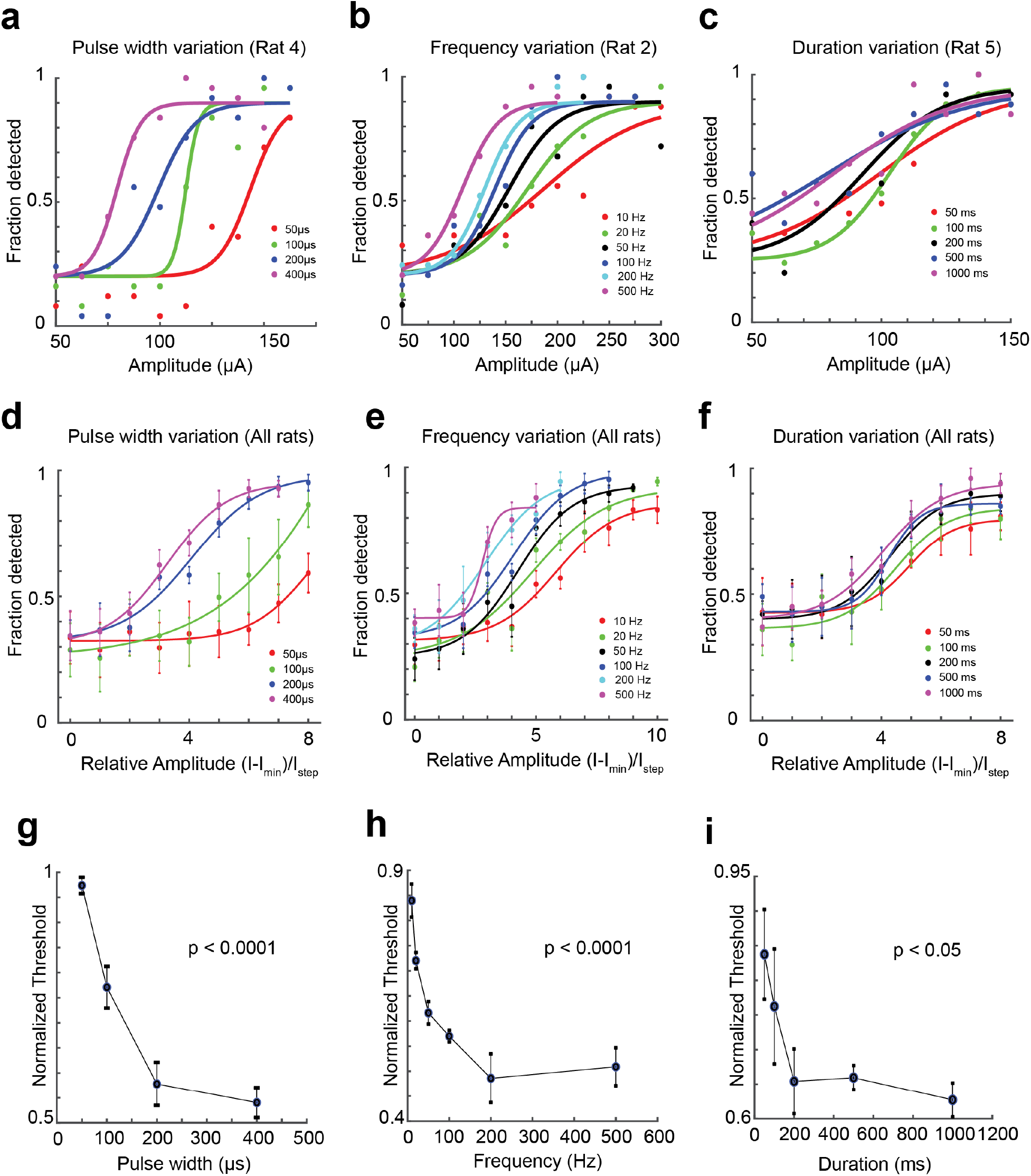
Psychophysical evaluation of SCS sensory detection with varying stimulation parameters in rats. Once rats learned to detect SCS stimuli at the standard parameters (frequency: 100 Hz, pulse width: 200 μs, duration: 1 sec), we allocated different blocks of sessions where (pulse width & amplitude; panel ‘a’); (frequency & amplitude; panel ‘b’); and (duration & amplitude; panel ‘c’) were varied while keeping other parameters constant. a, b, c) Psychometric curves from representative rats showing a leftward shift of curves as the pulse-width (rat 4), frequency (rat 2), and duration (rat 5) of stimulation increased. d, e, f) Psychometric curves with averaged data across five rats indicate leftward shift as pulse-width, frequency, and duration are increased. X-axis represents relative amplitude values (for each rat raw amplitude values were subtracted by minimum amplitude and the difference was divided by amplitude step size). Circles and error bars are mean ± sem across five rats. Curves in panels ‘a-f’ are sigmoidal fits. Panels ‘g’, ‘h’, and ‘i’ indicate normalized detection thresholds. Thresholds were calculated as 75% fraction detected at different stimulation parameters consistent with panels ‘a-f’ and then normalized by maximum amplitude used in the experimental block of each rat. Circles and error bars are mean ± sem across five rats. P-values were calculated using repeated measures one-way ANOVA.

### Sensory discrimination in primates

We then trained monkeys K and O to discriminate SCS that varied in frequency as well as location of stimulation. On frequency discrimination (100 Hz vs 200 Hz, Figure 6a), monkey O’s performance improved from 29% on day 1 to 96% on day 17 of training, while monkey K’s performance improved from 74% on day 1 to 81% on day 11 (Figure 6b). Spatial discrimination was achieved by stimulating electrode pair 1 versus electrode pair 2 (Figure 6c), where monkey O’s performance improved from 46% on day 1 to 97% on day 11 and monkey K’s performance improved from 74% on day 1 to 86% on day 7 (Figure 6d).

**Figure 6:**
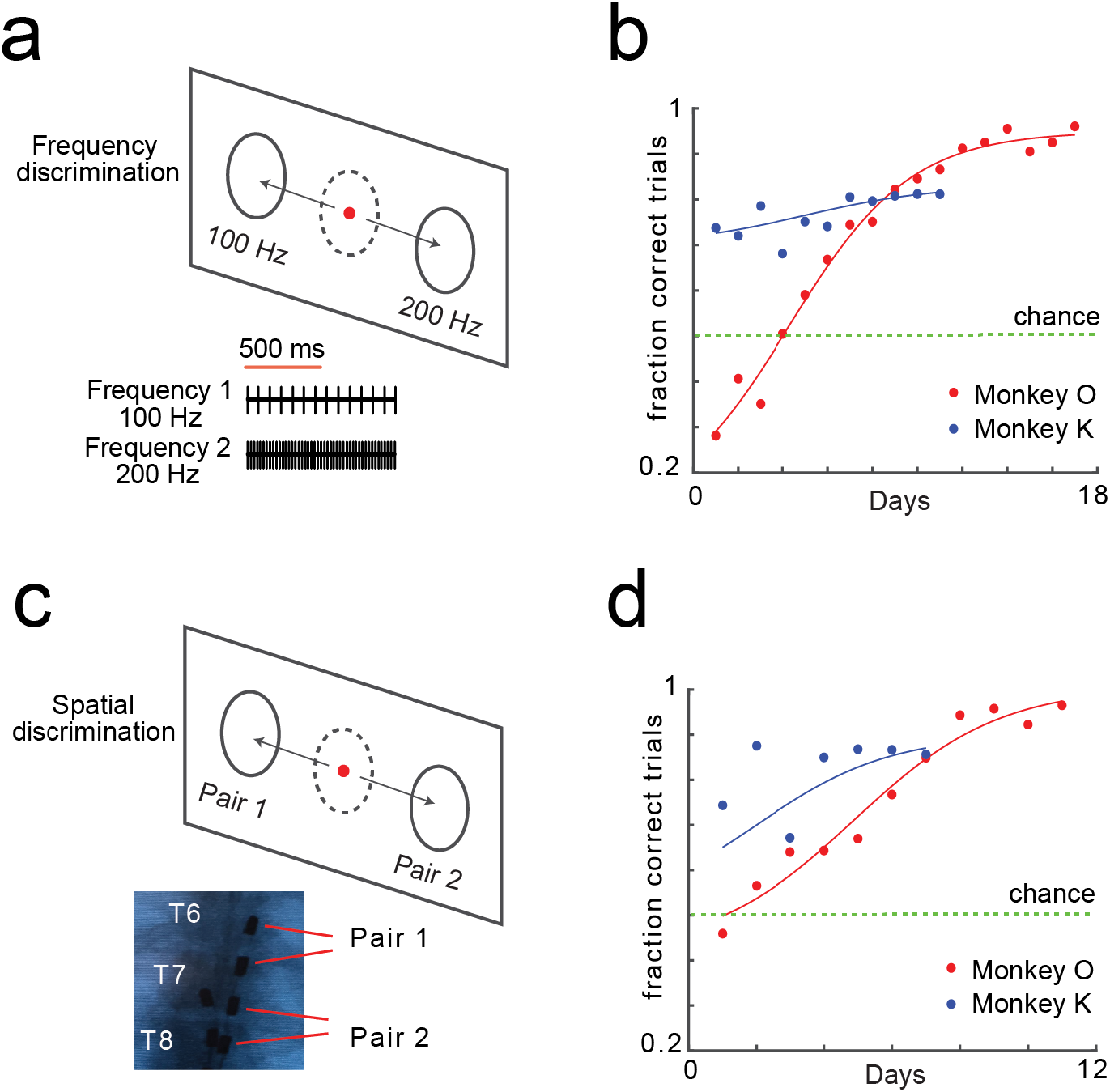
Monkeys learned to discriminate SCS stimuli varying in frequency and spatial location of electrodes. a,b) Monkeys K and O learned to discriminate SCS stimuli delivered at the same electrode location but varying in frequency (100 Hz vs 200 Hz). Monkey O was stimulated at same amplitude while monkey K was stimulated at the respective threshold amplitude for 100 Hz and 200 Hz. c,d) Monkeys K and O learned to discriminate stimulation delivered at electrode pair 1 (T6 - T7 spinal level) vs electrode pair 2 (T7-T8 spinal level). Curves in panels b and d indicate sigmoidal fits to fraction correct trials displayed as a function of training days.

### Sensory discrimination and Weber’s law in rats

We had previously shown that rats can learn to discriminate up to four different time-varying patterns of stimulation (14). In our current work, we trained rats to discriminate SCS stimuli with different frequencies using the same behavioral setup that was used for the detection task (Supplementary Figure 4a). In the basic training, rats learned to discriminate between 10 Hz and 100 Hz of stimulation delivered at pulse-width of 200 μsec, and duration of 1 sec (Supplementary Figure 4b). Thereafter, we studied whether discrimination of sensations induced by different stimulation frequencies follows the rules of Weber’s law (16), which states that Just-Noticeable Differences (JNDs) between a standard frequency and comparison frequency should linearly increase with the standard frequency of stimulation. To this end, we determined JNDs at different standard frequencies (10 – 100 Hz) where the comparison frequency was higher than the standard (Figure 7a). We observed that JNDs had a significant linearly increasing relationship with the standard frequency of stimulation (Figure 7b, p< 0.0001, linear regression test). After that, we kept the standard as a higher frequency value (100 – 400 Hz) and decreased the comparison frequency randomly from that value (Figure 7c). In this case also, we observed that the JNDs for lower frequency comparison significantly increased linearly as the standard frequency increased (Figure 7d, p< 0.0001, linear regression test). These results suggest that the JND rule defined by Weber’s law holds true for sensory discrimination of SCS frequencies.

**Figure 7:**
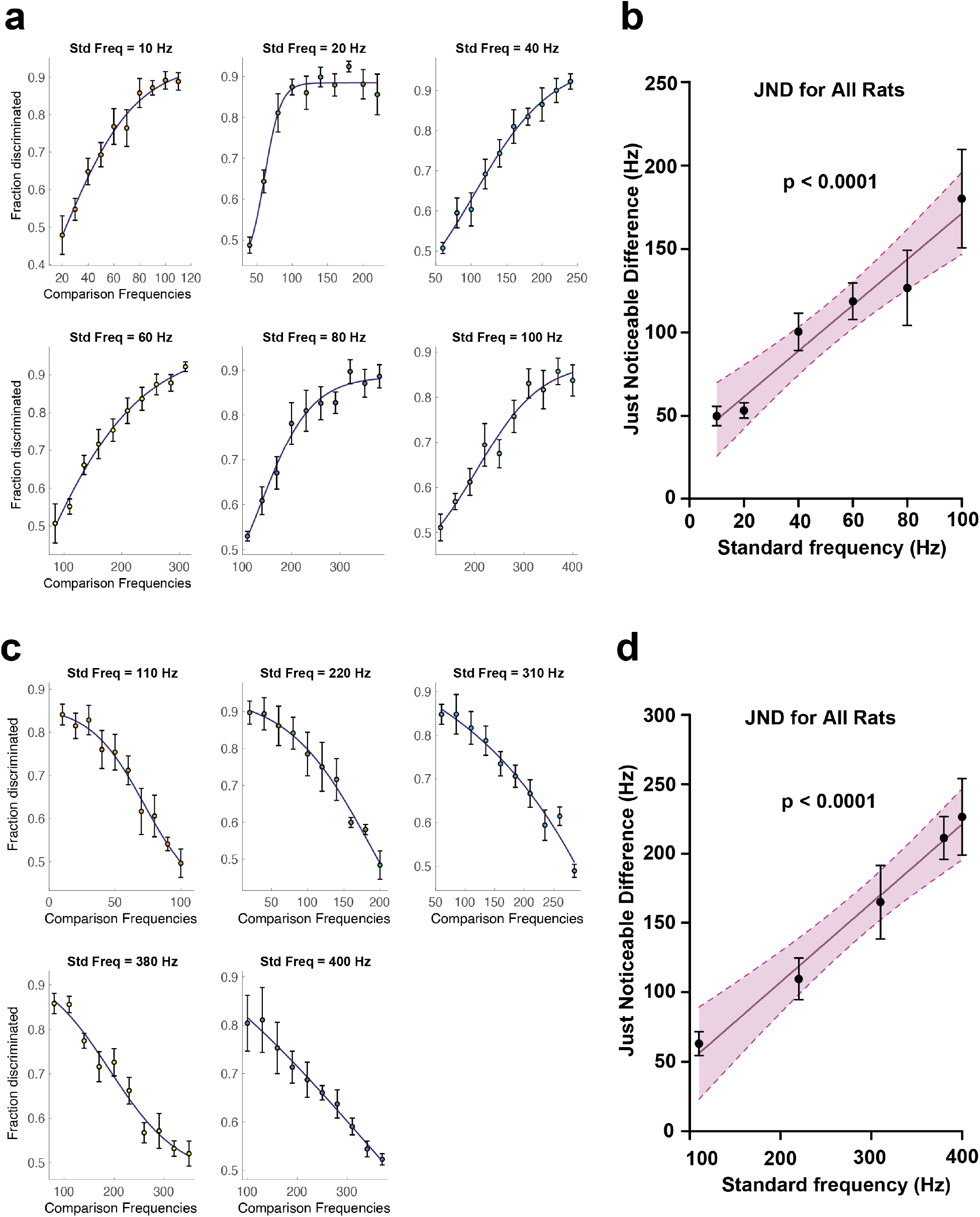
Discrimination of higher and lower comparison frequency obeys Weber’s Law. a) Fraction of trials correctly discriminated from standard frequency of 10 Hz, 20 Hz, 40 Hz, 60 Hz, 80 Hz, and 100 Hz, when compared to a range of higher frequency stimuli. c) Fraction of trials correctly discriminated from standard frequency of 110 Hz, 220 Hz, 310 Hz, 380 Hz, and 400 Hz, when compared to a range of lower frequency stimuli. Circles and error bars in ‘a’ and ‘c’ indicate mean ± sem. Curves indicate sigmoid fits to the data. Just noticeable difference (JND) is considered as the comparison frequency value that achieves 75 % discrimination ability. b, d) Just noticeable difference as a function of standard frequency is indicated by black circles and error bars. Black line is linear regression fit to the data and pink bounds indicate 95% confidence bound of the regression line. JNDs associated with higher frequency and lower frequency comparison were significantly linearly related with standard frequency. P-values were calculated using linear regression test.

## Discussion

In this study, we found that rhesus monkeys and rats can learn to detect and discriminate artificial sensations induced by SCS following several days of exposure. The threshold for detecting SCS decreases with increasing pulse width, frequency, and duration of the stimulus. We also documented the ability of monkeys to discriminate sensations that are generated by stimulation pulses with varying frequency and spatial location. In rats, we showed that the just-noticeable differences (JNDs) from a perceivable stimulus frequency were linearly related to the perceivable frequency when it was compared with a stimulus that had either higher or lower frequency. These results demonstrated the unique ability of SCS as a novel transmission channel to the brain to encode highly contextual sensory information.

Our results on behavioral sensitivity to detection of sensation in rodents and primates were comparable to those observed with Intracortical Microstimulation (ICMS) of S1 (5, 17). Notably, sensitivity to stimulus amplitude increased with increasing pulse width, frequency, and duration of stimulation. ICMS amplitude using currently FDA-approved UTAH arrays is usually restricted below ∼100 µA due to the possibility of brain tissue damage, which limits the amplitude range for neuroprosthetic applications between detection thresholds of 20 - 50 µA and maximum allowable safe amplitude of ∼100 µA. In our SCS study, detection thresholds had a wider range from 200 – 800 µA at different stimulation settings and all detection thresholds were consistently on the non-damaging side of the boundary between damaging and non-damaging stimulation delineated by Shannon equation [log(D) = k – log (Q), with k = 1.85] on log charge density versus log charge per phase plot (Supplementary Figure 3a) (18). Assuming a commercially accepted maximum charge density of 30 µC/cm^2^ and minimum pulse-width of 50 µs (19), it could be estimated that maximum SCS current of ∼80 mA in monkeys and ∼3 mA in rats could be delivered using electrode contacts (monkeys: 0.1319 cm^2^; rats: 0.005 cm^2^) reported in our study without causing tissue damage.

Frequency modulation has been historically considered a promising method for providing sensory feedback with several studies showing that animals are capable of discriminating ICMS frequencies and that frequency modulation obeys Weber’s law. (4, 20-22), While ICMS amplitude modulation in monkeys failed to follow Weber’s law (5), experiments in rats showed that modulation of perceived intensity by amplitude and pulse-width modulation followed Weber’s law (17). In our study, we investigated whether rats and monkeys could learn to discriminate SCS frequencies. Although monkeys O and K learned to discriminate 100 Hz SCS from 200 Hz, after taking a closer look at their learning curves it was evident that they displayed different learning behaviors in the frequency discrimination task (Figures 6b and 6d). Monkey O started at lower discrimination performance at earlier training sessions but reached higher level of performance toward the end of training, whereas monkey K’s performance started higher than chance and improved marginally as the training progressed. These differences in learning behavior could be attributed to the fact that monkey O was stimulated at constant amplitude for both frequency values (100 Hz and 200 Hz) while monkey K was stimulated at each frequency’s threshold amplitude. It is quite possible that monkey O was discriminating differences in perceived magnitudes of the sensory percepts while monkey K was discriminating differences in the qualitative nature of the percepts induced. Rats are capable of discriminating temporal patterns of SCS when the number of pulses are kept constant but the frequency of stimulus is varied (14). In addition to that, our current results indicate that JNDs associated with SCS frequency discrimination in rats clearly follow Weber’s law because JNDs linearly increased with standard stimulation frequency (Figure 7b and 7d). There were at least three discriminable percepts between 10 Hz and 200 Hz. It can be argued that applying frequency modulation simultaneously with amplitude and pulse-width modulation would potentially increase the number of distinct discriminable percepts that are possible within the amplitude range allowed on current SCS electrodes. Therefore, further experiments exploring the relationship between sensory discrimination and frequency, pulse-width, amplitude, and duration of stimulation are necessary to understand how these parameters relate to perceived intensity and quality of sensation evoked by SCS.

A major advantage of SCS is its ability to target multiple dermatomes simultaneously with a single electrode array. In particular, a single, commercially available SCS lead with multiple contacts can evoke sensations in multiple dermatomes simultaneously due to the bilateral sensory representation of the entire lower body in the ascending dorsal column fibers. It is quite apparent from our results that monkeys can learn to discriminate the spatial location of the sensations evoked by SCS (Figure 6d). This suggests that we can take full advantage of the medio-lateral/rostro-caudal somatotopy represented in the dorsal column fibers in combination with spatiotemporal stimulation patterns to electrically induce targeted tactile or proprioceptive sensations in the body. This view is also supported by evidence from computational studies which indicate that epidural SCS activates dorsal column fibers up to a depth of 0.2-0.25 mm from the dorsal surface (23-27). However, additional work on miniaturizing electrode contacts and accurate mapping of SCS-induced sensations needs to be performed to be able to elicit precise sensations. In addition, the ability of SCS to modulate neuronal activity in supraspinal brain structures is quite desirable from a neuroprosthetic as well as a therapeutic application standpoint (14, 28-31). A major limitation of our study is the short experimental time (Supplementary Figure 1a) we had available for primates – maximum of 5 months – due to the risk of infection associated with externalized SCS leads. A fully implantable stimulation system, like the one implanted in chronic pain patients could potentially extend our study indefinitely and allow us to perform longer experiments in monkeys. Nevertheless, in rodents we were able to perform longer post-implant experiments because the electrodes and their wires were fully enclosed inside the body.

In conclusion, we have successfully demonstrated that SCS can be used to encode sensory information in both rats and monkeys and all together our results demonstrate that SCS-induced sensations obey a robust encoding scheme in the brain. Additionally, our behavioral experiments serve as a test bed for future animal studies which could elucidate the neural mechanisms underlying SCS-based sensory detection and discrimination. We envision that SCS can be developed as an artificial sensory feedback channel for delivering targeted tactile and proprioceptive information to the brain.

## Methods

All animal procedures were approved by the Duke University Institutional Animal Care and Use Committee and in accordance with National Institute of Health Guide for the Care and Use of Laboratory Animals. Three adult rhesus macaque monkeys (Macaca mulatta), monkeys ‘M’, ‘O’, and ‘K’ and Long Evans rats (300 -350 g) participated in the experiments.

### Monkey spinal implant surgery

Monkeys M, K, and O were implanted with 8-contact cylindrical percutaneous leads (Model 3186, diameter 1.4mm, contact length 3 mm, spacing 4 mm, Abbott Laboratories) bilaterally to the spinal mid-line in the dorsal epidural space at T6 – T8 spinal level. Monkey M had two leads with 8 stimulation contacts each, monkey O had two leads, one with 8 contacts (right side) and one with 4 contacts (left side, Model 3146, similar electrode contact dimensions as Model 3186), and monkey K had two leads with 8 stimulation contacts each. Experimental procedures for monkeys M, O, and K lasted approximately 45, 135, and 150 days post-implant after which the leads were explanted (Supplementary Figure 1a).

Implant surgery was performed under general anesthesia using standard procedures typical of human implantation (for details see supplementary information and Supplementary Figures 1b and 1c). Once leads were implanted in the epidural space, a small hole was created in the skin off-midline to externalize the distal end of the lead. Externalized leads were enclosed in a custom plastic cap which was sutured to the skin (Supplementary Figure 1d). The plastic cap allowed for access to the leads by the researcher but protection from the animal. The animal wore a protective vest after surgery and throughout the experimental period which prevented its access to the plastic cap sutures to its back.

### Monkey SCS detection task

Monkeys were trained to perform a two-alternative forced choice task where they were seated in a chair facing a computer monitor which indicated trial progression (Figure 1b). On each trial a center target appeared, and the monkey had to move a cursor which was joystick-controlled inside the center target (Figure 1c). After a brief hold period of 500 milliseconds (ms) inside the center target, two targets appeared on either side of the center target. Each monkey had to hold the cursor inside the central target for a brief period of 500 ms – 1 second. During this hold period movement cues were presented. If SCS was presented (charge-balanced, cathode-first, 200 µs biphasic square pulses at 100 Hz for 1 second using custom microstimulator (32)), then the monkey had to move the cursor to the left target to obtain a juice reward. If SCS was absent, then the monkey had to move the cursor inside the right target. At the end of the hold period, the center target disappeared, thus cuing the monkey to initiate cursor movement toward the reward. Incorrect target reaches were not rewarded. Initiation of movement prior to the end of the hold period terminated the trial without reward and a blank screen was displayed 3 seconds before starting next trial. The learning performance of monkeys was studied using percentage correct (PC) trials.

### Monkey psychometric evaluation of detection thresholds

Once monkeys were trained on the detection task, detection thresholds were determined for different electrode combinations using psychometric testing. Particularly, during ‘Stimulation ON’ i.e. ‘left target rewarded’ trials, the stimulation amplitude was randomly varied from 50 μA to a manually determined upper limit which was below the motor threshold. Only left target trials were analyzed and a percentage correct (PC) performance at each stimulation amplitude value was determined. A sigmoid curve was fit to the PC values and 75% was considered as detection threshold. This was repeated for several electrode combinations.

### Monkey electrode mapping

In monkey K, once sensory detection thresholds were determined, we mapped the location of electrode pairs to location on the monkey’s body by sedating the monkey and stimulating those electrodes above the sensory threshold values. Areas on the body surface that elicited minor muscle twitches or skin flutter were marked (Figure 2c, left) and the motor thresholds were noted. In monkey O, these observations were not made under sedation but while it was seated in the primate chair.

### Monkey detection thresholds as stimulation parameters vary

During sets of consecutive sessions, we varied amplitude and frequency or amplitude and pulse-width or amplitude and duration of stimulation while keeping other parameters constant (for stimulation parameter ranges, see Supplementary Table 1). The standard parameters that remained constant while others were varied were frequency: 100 Hz, pulse width: 200 μsec, and duration: 1 sec. In monkey K, we varied frequency (10 – 1000 Hz) and duration (1 – 2000 ms) of stimulation simultaneously while keeping pulse width and amplitude constant at 200 μs and 325 μA respectively. Detection thresholds for each monkey were normalized by maximum amplitude used in the experiment block of that monkey before statistical analysis.

### Monkey sensory discrimination

In the frequency discrimination task, each monkey was instructed to move the cursor inside the left target for 100 Hz and right target for 200 Hz respectively (Figure 6a). In the spatial discrimination task, the monkey was instructed to move the cursor inside the left target for electrode pair 1 and inside right target for electrode pair 2 respectively (Figure 6c).

### Rat pre-training and SCS electrode implantation

Moderately water deprived rats were placed inside the behavioral chamber for 2 days to acclimatize to the behavioral environment. The behavioral chamber had two doors on one side of the walls which enclosed water reward ports (Figure 4a), slightly modified from the one previously described (33). Rats were gradually trained to receive water reward from the ports. Initially, both reward doors were kept open and rats learned to receive water by licking at the water dispensing spout. Later, the doors were kept closed and would open a few seconds after the house light turned on. In subsequent sessions, left and right doors would open on alternate trials and rats learned to obtain reward from each port alternatively. The pre-surgical training period lasted approximately 8-10 days.

Thereafter custom designed SCS electrodes (1 mm X 0.5 mm contacts arranged transversely in a bipolar configuration with 0.25 mm spacing using a 0.025mm thickness platinum foil, Goodfellow Cambridge Limited, England) were implanted into the epidural space under T3 vertebra as described in our previous article (29). After the rats recovered from the spinal surgical procedures, cathode leading stimulation pulse trains were delivered at the SCS electrodes using a multi-channel constant current stimulator (Master-8, A.M.P.I, Jerusalem, Israel) at stimulation settings which were determined depending on the behavioral task under consideration (Figure 4a).

### Rat sensory detection task

After recovery from surgery, rats were introduced to a two-alternative forced choice task (2AFC) to learn detection of SCS stimuli in the chamber (Figure 4b). At the beginning of each trial, a light in the chamber was turned on for 1 second as a reminder for the animals to pay attention. After the light turned off, the rats received a sensory cue for 1 second. The sensory cue either consisted of cathode-leading bipolar square pulse trains (pulse width: 200 μs, Frequency: 100 Hz, duration: 1 s) or no stimulation pulses (interval: 1 s) at each trial. After a brief delay of 0.5 seconds both reward doors opened, and rats had to respond by choosing the left door for ‘SCS ON’ trials and right door for ‘SCS OFF’ trials. Incorrect responses were not rewarded. During the learning of this basic detection task, the intensity of the delivered current was determined before each session and set using procedures described before (14, 29, 31) (mean±std, intensity at 100 Hz was 243.7 ± 57.9μA).

### Rat sensory detection psychophysics

Once rats learned the basic sensory detection task and their performance was above 80%, stimulation parameters were varied in a systematic manner during the SCS-ON trials. In different experimental sessions, stimulation parameters such as frequency and amplitude, pulse-width and amplitude, and pulse train duration and amplitude were varied while other parameters were kept constant (standard parameters: pulse width: 200 μs; Frequency: 100 Hz; duration: 1 s), and sensory detection threshold amplitude was determined (for stimulation parameter ranges see Supplementary Table 1). For all the conditions, the detectable level of the amplitude was defined as 75% accuracy of behavioral performance. Detection thresholds for each rat were normalized by maximum amplitude used in the experiment block of that rat before statistical analysis.

### Rat sensory discrimination task

In a 2AFC task, rats were presented with either a low frequency stimulus or a high frequency stimulus during the sensory cue period in the behavioral chamber. For either frequency, the stimulus was delivered at the same amplitude (determined at each session), pulse width (200 μs), and duration (1 sec). After a brief delay period following sensory cue presentation, rats had to choose the left door for higher frequency stimulus and right door for lower frequency stimulus to obtain reward (Supplementary Figure 4a). Initially, rats were trained to discriminate between 10 Hz and 100 Hz frequency. Incorrect trails were not rewarded.

### Weber’s law and sensory discrimination

Once rats learned to discriminate 10 Hz stimulus from 100 Hz stimulus, demonstrated by consistent discrimination performance above 80%, the lower frequency (standard frequency) was kept constant during right door trials while the higher frequency (comparison frequency) was randomized between 20 Hz and 110 Hz during left door trials. Sensory discrimination performance between a standard frequency and comparison frequency was determined as the fraction of trials successfully discriminated for that particular standard and comparison pair. Thus, fraction trials discriminated was plotted as a function of comparison frequency to obtain Just-Noticeable Differences (JNDs) determined as 75 % value on the curve. The same experiment was repeated for different standard frequency values such as 20 Hz vs (40 – 220 Hz), 40 Hz vs (60 – 240 Hz), 60 Hz vs (85 – 310 Hz), 80 Hz vs (110 – 380 Hz), and 100 Hz vs (130 – 400 Hz) to obtain JNDs for each standard frequency value and to test for Weber’s law.

Similarly, the experiment was repeated for discrimination of a low-frequency comparison stimulus from a high-frequency standard stimulus. The standard frequency values were 110, 220, 310, 380, and 400 Hz, while the comparison frequency was randomized from (100 – 10 Hz), (200 – 20 Hz), (285 – 60 Hz), (350 – 80 Hz), and (370 – 100 Hz) respectively for each of the standard frequency values. JNDs were calculated for each of the standard frequency values as explained before to test for Weber’s law.

## Statistical Analysis

Repeated measures one-way ANOVA was used to test the significance of relationship between stimulation parameters and sensory detection thresholds in both rats and monkeys (Figures 3h, 3i, 3j, 5g, 5h, and 5i). To test whether JNDs were significantly linearly related to standard frequency in the sensory discrimination task, the linear regression test was used (Figures 7b and 7d).

## Acknowledgements

We thank Gary Lehew for assistance with experimental setup, Tamara Phillips for assistance with monkey handling, Paul Thompson for assistance with primate task software, Laura Oliveira and Susan Halkiotis for technical assistance, and Joseph O’Doherty for comments on previous version of manuscript. This research was supported by Duke Institute for Brain Sciences Germinator Award and Duke Neurosurgery Research Support offered to Amol Yadav, NIH R25 awarded to Max Krucoff, and Hartwell Foundation grant awarded to Miguel Nicolelis.

## Author Contributions

A.P.Y., S.L., M.O.K., and M.A.L, designed experiments, A.P.Y. and S.L. performed rodent experiments and surgeries, M.O.K and M.M.A performed primate surgeries, A.P.Y performed primate experiments, A.P.Y. and S.L. analyzed data, A.P.Y. drafted manuscript, all authors edited manuscript, M.A.L.N. supervised work.

## Supplementary Information and Figures

### Primate implant surgery and timeline

Two weeks prior to surgery, the animal was acclimated to a protective vest. One day before surgery, the animal’s back was shaved with an electrical clipper under mild sedation. The day of surgery, the animal underwent general anesthesia and endotracheal tube placement by staff veterinarians. The animal was then positioned prone on a radiolucent fluoroscopy table. The spine was placed into gentle flexion to open the interspinous spaces by placing the knees and pillows under the abdomen of the monkey. Pressure points were carefully padded. The surgical area was cleaned and prepped with 2% chlorhexidine gluconate and 70% isopropyl alcohol solution (ChloraPrepTM) and allowed to dry for 3 minutes. The surgical area was then draped in a sterile fashion. A dose of antibiotics was given within one hour prior to incision.

An anterior-posterior (AP) C-arm fluoroscopic x-ray was then taken to mark the level of the lowest rib, and the radiographic shadows of the spinous processes were positioned in the middle of the pedicles to ensure minimal rotation. After injecting a local anesthetic (1.5% lidocaine with 1:100,000 epinephrine), a 2cm incision was made at this location down to the fascia, and a self-retaining retractor was placed to hold the skin open. A trocar was then introduced through the fascia just lateral to midline between the spinous processes and advanced so the tip just entered the epidural space. Epidural location was confirmed using a loss of resistance technique, fluoroscopic x-ray, and lack of cerebrospinal fluid (CSF) (34). Next, an 8-contact cylindrical percutaneous lead was introduced through the trocar and steered through dorsal epidural space to a target location in the mid-thoracic spine, or approximately across T8 (Supplementary Figure 1b). Once in a satisfactory position, the stylet and trocar were removed. The lead was then secured to the fascia using a 2-0 silk suture, and then a subcutaneous suprafascial pocket was created off midline to secure lead excess. A small hole was then created in the skin off midline to externalize the distal end of the lead. The same procedure was then repeated on the opposite site, ultimately positioning the electrodes next to one another in the midline dorsal epidural thoracic space with a slight cranio-caudal offset (Supplementary Figure 1c).

Once both electrodes were externalized, the incision was closed with inverted dermal vicryl sutures and a running 4-0 nylon suture. The incision was then covered with a thin layer of bacitracin ointment, and piece of telfa was then placed over the incision and stapled to the skin. Next, a custom plastic cap was sutured to the animal’s skin with 3-0 nylon sutures to cover the externalized leads (Supplementary Figure 1d). Such a cap allowed for access to the leads by the researcher but protection from the animal. The telfa was removed 3 days after surgery. Surgical sutures were removed 7-10 days after surgery. The sutures in the cap needed to be replaced approximately monthly under sedation.

Monkey M’s right lead stopped working a few days post-implant while the left lead stopped working approximately 45 days post-implant. Hence the leads were explanted thereafter. Monkey O’s left lead stopped working a few days post-implant while the right lead worked for 135 days after which they were extracted due to a skin infection at the externalization site. Monkey K’s both left and right leads were working until they were explanted approximately 150 days after surgery due to a skin erosion at externalization site.

**Supplementary Figure 1:**
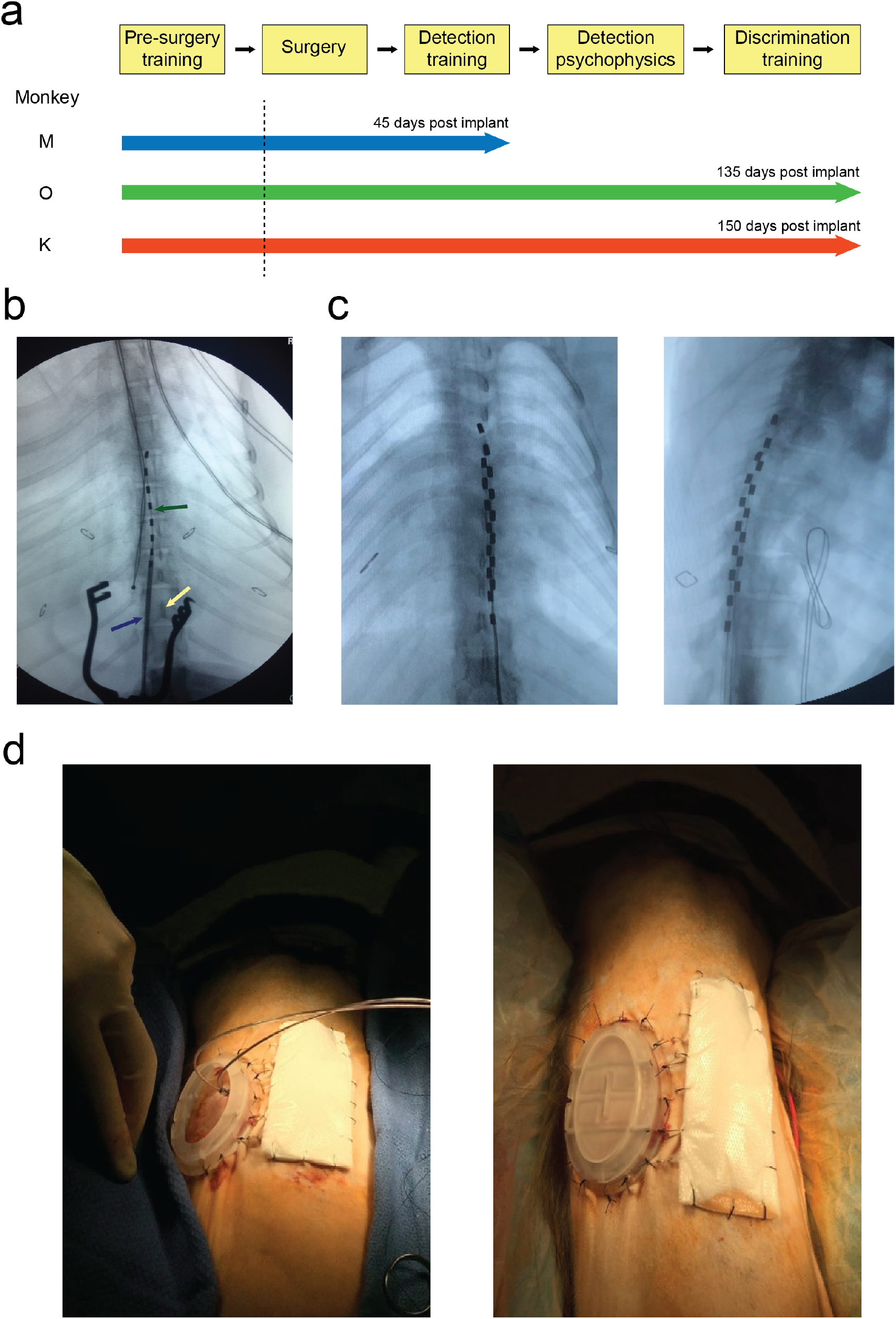
Timeline of primate experiments and primate surgery procedures. a) Monkeys were trained on the joystick task prior to spinal implant surgery. After they recovered from the surgery behavioral training began for detection of SCS-based sensations. After that sensory detection psychophysical experiments were performed followed by training on discrimination of SCS-based sensations. Monkey M learned the SCS detection task, however, 45 days after surgery its implanted spinal electrodes stopped working and hence were explanted. Spinal implants of Monkey O and K lasted until they both were trained on the SCS sensory discrimination task. However, 135 and 150 days after surgery for both monkeys the implants stopped working and were subsequently explanted. b) Anterior-posterior (AP) fluoroscopic x-ray showing the introduction of the 8-contact cylindrical lead through the trocar into the mid-thoracic dorsal epidural space. Blue arrow: trocar, Green arrow: 8-contact electrode array, Orange arrow: spinous process. c) Final AP (left) and lateral (right) x-rays showing placement of dorsal mid-thoracic epidural electrodes with slight cranio-caudal offset. d) Externalized electrodes with custom plastic cap sutured to the skin, and incision temporarily covered with telfa stapled to the skin. Cap open (left), cap closed (right).

**Supplementary Figure 2:**
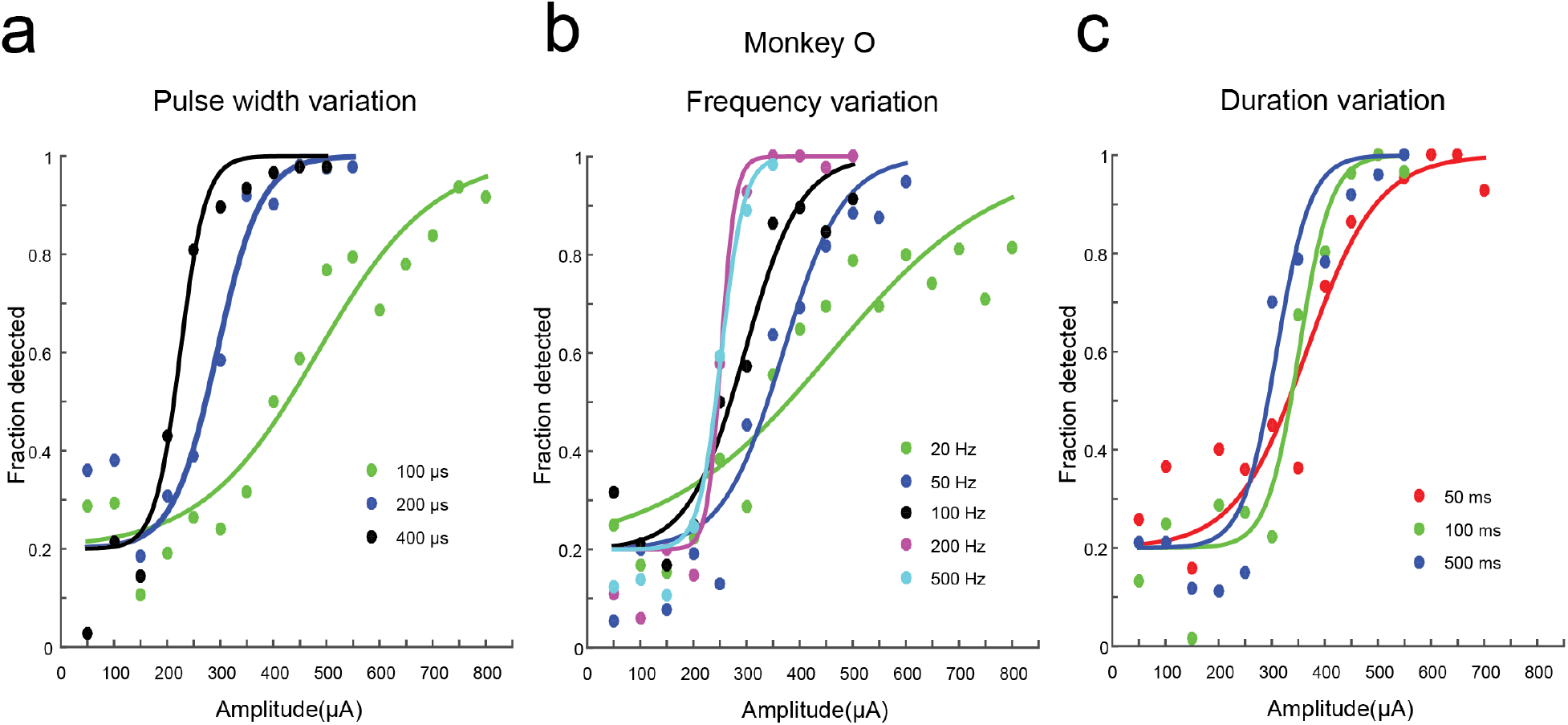
Psychophysical evaluation of SCS sensory detection in monkey ‘O’ with varying stimulation parameters. Once monkey O learned to detect SCS stimuli at the standard parameters (frequency: 100 Hz, pulse width: 200 μs, duration: 1 sec), we allocated different blocks of sessions where (pulse width & amplitude; panel ‘a’); (frequency & amplitude; panel ‘b’); and (duration & amplitude; panel ‘c’) were varied while keeping other parameters constant. Psychometric curves in a-c are sigmoidal fits to fraction of trials that were successfully detected as a function of amplitude.

**Supplementary Figure 3:**
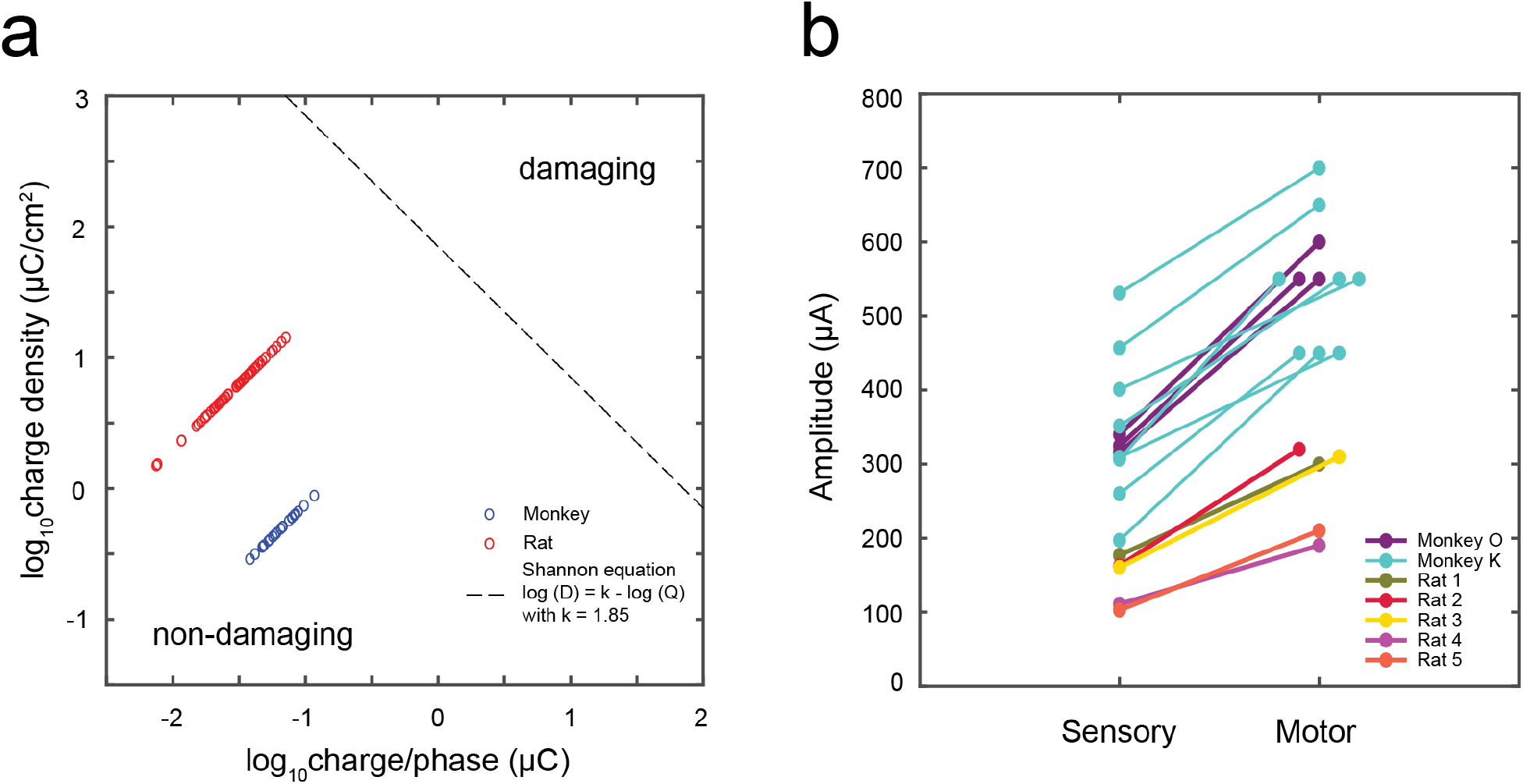
Charge versus charge density and sensory versus motor thresholds. a) Sensory thresholds for monkeys (blue circles) and rats (red circles) determined by psychophysical evaluation at multiple stimulation parameters of pulse-width, frequency, and duration were below tissue damaging levels. Black dotted line indicates boundary between damaging and non-damaging stimulation defined by k = 1.85 in the Shannon equation [log(D) = k – log (Q)] on a log charge density (D) versus log charge per phase (Q) plot. b) Sensory thresholds were considerably lower than the amplitude levels where muscle twitches or skin fluctuations were observed on the animals’ body. Purple and turquoise circles represent multiple electrode pairs tested in monkeys O and K respectively.

**Supplementary Figure 4:**
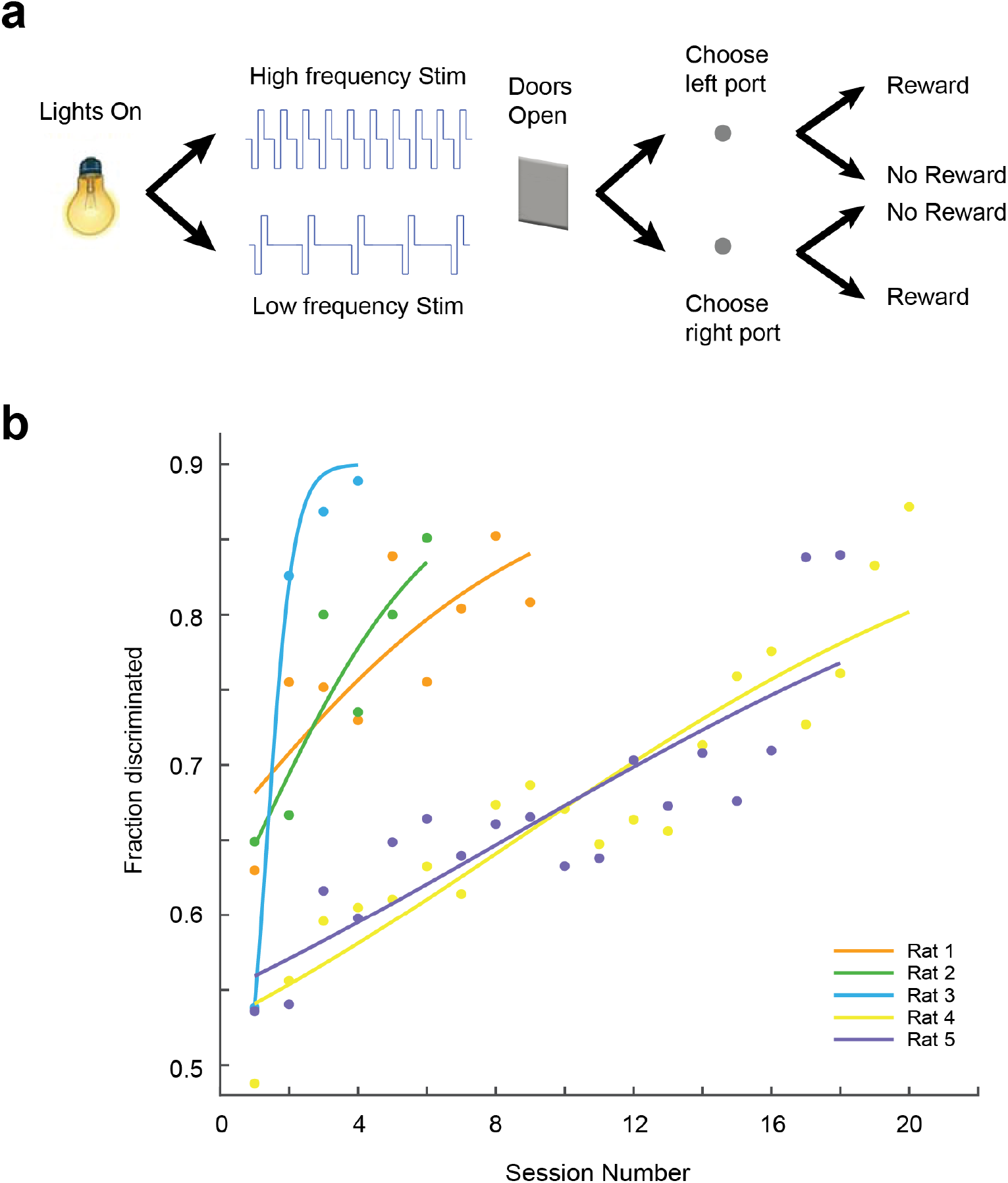
Rats learned to discriminate SCS stimuli varying in frequency. a) Behavioral setup for frequency discrimination task. Rats had to choose left reward port for higher frequency of SCS and right reward port for lower frequency of stimulation. Stimulation was delivered at a constant amplitude at the standard parameters (pulse width: 200 μs, duration: 1 sec) and rats had to choose the correct reward port associated with each frequency to obtain water reward. b) Learning curves for five rats that learned to discriminate two frequencies (10 Hz vs 100 Hz). Curves indicate sigmoidal fits to fraction of trials that were successfully discriminated as a function of session number.

**Supplementary Table 1:** Stimulation parameter ranges tested at psychophysical evaluation of sensory detection. Animals were initially trained to detect SCS stimuli at standard stimulation parameters – frequency: 100 Hz, pulse-width: 200 µs, and duration: 1000 ms) as shown in Figures 2a and 4c. After that sensory thresholds were determined by using stimulation settings shown above. In blocks of consecutive sessions, amplitude and frequency or amplitude and pulse-width or amplitude and duration of stimulation were varied while keeping other parameters constant. In additional block of sessions in monkey K, frequency (10 – 1000 Hz) and duration (1 – 2000 ms) of stimulation were simultaneously varied while keeping pulse width and amplitude constant at 200 μs and 325 μA respectively.

| Animal | Amplitude | Pulse Width | Frequency | Duration |
| --- | --- | --- | --- | --- |
| Monkey O | 50 – 800 μA, 50 μA step | 100 – 400 μs | 20 – 500 Hz | 50 – 500 ms |
| Monkey K | 50 – 800 μA, 50 μA step | 50 – 400 μs | 10 – 500 Hz | 50 – 1000 ms |
| Rat 1 | 50 – 325 μA, 25 μA step | 50 – 400 μs | 10 – 500 Hz | 50 – 1000 ms |
| Rat 2 | 50 – 325 μA, 25 μA step | 50 – 400 μs | 10 – 500 Hz | 50 – 1000 ms |
| Rat 3 | 50 – 325 μA, 25 μA step | 50 – 400 μs | 10 – 500 Hz | 50 – 1000 ms |
| Rat 4 | 50 – 187.5 μA, 12.5 μA step | 50 – 400 μs | 10 – 500 Hz | 50 – 1000 ms |
| Rat 5 | 50 – 175 μA, 12.5 μA step | 50 – 400 μs | 10 – 500 Hz | 50 – 1000 ms |

